# Electric Field Induced Wetting of a Hydrophobic Gate in a Model Nanopore Based on the 5-HT_3_ Receptor Channel

**DOI:** 10.1101/2020.05.25.114157

**Authors:** Gianni Klesse, Stephen J. Tucker, Mark S.P. Sansom

## Abstract

In this study we examined the influence of a transmembrane voltage on the hydrophobic gating of nanopores using molecular dynamics simulations. We observed electric field induced wetting of a hydrophobic gate in a biologically inspired model nanopore based on the 5-HT_3_ receptor in its closed state, with a field of at least ∼100 mV nm^−1^ was required to hydrate the pore. We also found an unequal distribution of charged residues can generate an electric field intrinsic to the nanopore which, depending on its orientation, can alter the effect of the external field, thus making the wetting response asymmetric. This wetting response could be described by a simple model based on water surface tension, the volumetric energy contribution of the electric field, and the influence of charged amino acids lining the pore. Finally, the electric field response was used to determine time constants characterising the phase transitions of water confined within the nanopore, revealing liquid-vapour oscillations on a time scale of ~5 ns. This time scale was largely independent of the water model employed and was similar for different sized pores representative of the open and closed states of the pore. Furthermore, our finding that the threshold voltage required for hydrating a hydrophobic gate depends on the orientation of the electric field provides an attractive perspective for the design of rectifying artificial nanopores.

**ToC/Abstract Graphic:** 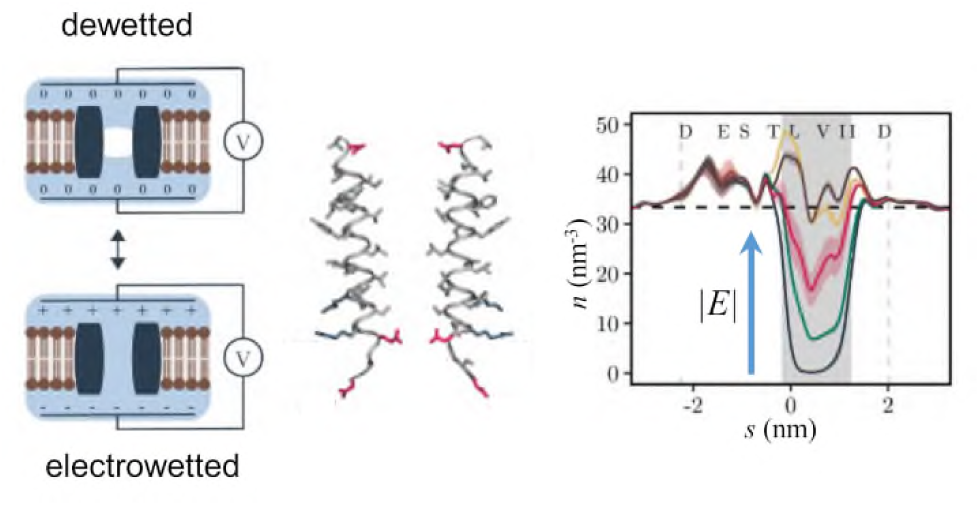

## Introduction

Nanopores enable the permeation of water and small molecules through membranes that separate aqueous compartments. The size of these permeant molecules is typically comparable to the diameter of the pores involved (≤ 1nm). Consequently, any interaction of the permeant molecules with the lining of the pores will determine the functional properties of the permeation process ^1–4^. In biological membranes, ion channels are structurally dynamic protein nanopores that switch between functionally open and closed conformations to control the rapid permeation of ions and water across the membrane ^5–6^. However, when open, a typical channel pore has an internal radius of ~0.5 nm and a length of ~5 nm that creates a nano-confined environment where the precise shape and related dynamic physicochemical properties of the pore will also influence the complex behaviour of water and of ions, and therefore their permeation across the membrane. For example, we have previously shown how ion permeation can be influenced by both the pore radius and local hydrophobicity of the pore lining ^7–9^. In such cases, the presence of band of hydrophobic amino acid side chains lining a pore can induce a local liquid-to-vapour phase transition resulting in that section becoming devoid of liquid water. Such pore ‘de-wetting’ creates a free energy barrier for ion permeation. Thus, although ion permeation can occur through polar regions only just wider than the radius of the permeating ion, a hydrophobic region of comparable dimensions can prevent ion permeation without complete steric occlusion of the pore. This process has been referred to as hydrophobic gating and has enhanced our understanding of the mechanisms which control ion permeation through biological ion channels and synthetic nanopores ^10–13^.

The gating or regulation of hydrophobic gates in ion channels is thought to involve either structural changes in pore radii, or more subtle changes in hydrophobicity through e.g. rotation of helices that contain side chains of different polarity and examples of such hydrophobic gating mechanism have now been proposed in a variety of different channels ^12, 14–25^ and synthetic nanopores ^26–27^. However, nearly all biological membranes experience a potential difference of between 50 to 200 mV across them and molecular dynamics (MD) simulations of simple model nanopores show that application of a transmembrane electric field can hydrate an otherwise de-wetted hydrophobic constriction, which is thus rendered ion permeable ^28–29^. This electrowetting effect can be explained by incorporating the electrical energy of the pore volume element which is wetted into a simple thermodynamic model of gating (see Eq. 1 below; ^8^). In addition to this model, which assumes bulk water properties, a number of MD studies have explored the effects of an externally applied electric field on water molecules in nanoconfined hydrophobic environments ^4^. For example, simulations of water nanodroplets on hydrophobic surfaces ^30^ reveal that an electric field modifies the interfacial tension of a nanodroplet by influencing the orientation and hydrogen bonding structure of water molecules located on the droplet surface. Furthermore, MD simulations of water in planar hydrophobic confinement ^31^ also reveal that interfacial tensions decrease upon application of an electric field, alongside a field-induced change in the average number of hydrogen bonds formed by interfacial water molecules. Electrowetting has also been observed experimentally for several types of artificial biomimetic nanopores ^32^. For example it has been shown that hydrophobic nanopores in polyethylene terephthalate membranes can be wetted and functionally opened through the application of an electric field ^26^.

For biological ion channels, the role of electrowetting is even less well understood; most studies have been limited to MD simulations, demonstrating that the de-wetted hydrophobic gate of the MscS channel can hydrate when a transmembrane voltage of between 0.25 and 1.2 V is applied ^33–34^. Comparable electrowetting of a hydrophobic gate in a model protein nanopore has been simulated ^35^, and electrowetting has also been seen within the hydrophobic pores formed by carbon nanotubes ^36^. However, whilst it is clear that electrowetting can in principle open a hydrophobic gate, a detailed quantitative relationship between pore hydration probability and the applied electric field has notso far been examined for a biologically realistic model of an ion channel. It therefore remains unclear how such effects of electric fields depend on the structure of the pore and the strength of the applied field, or how robust such predictions are to variations in the precise type of water model used in the MD simulations. It is therefore important that these effects are captured by a theoretical model which may subsequently be used in a quantitative predictive fashion.

In this study we have examined the effects of an applied electric field on a hydrophobic gate within a model protein nanopore derived from the transmembrane pore of the 5-HT_3_ receptor, a biologically-relevant ion channel distributed throughout the central nervous system ^37^. Our results demonstrate that an electric field can induce wetting of the hydrophobic gate in this model nanopore, at values that that correspond to a (supra-physiological) transmembrane potential difference of ~1 V. A simple thermodynamic model can be used to quantitatively describe the increase in pore hydration probability with increasing electric field. Interestingly, the observed asymmetry of the hydration probability response curve can also be accounted for by the intrinsic electric field generated by the distribution of charged amino acid residues along the length of the nanopore which provides an attractive perspective for the design of intrinsically rectifying artificial nanopores.

## Results and Discussion

### A Model Nanopore based on the 5-HT_3_ Receptor

To quantify the effects of electrowetting of a hydrophobic gate, we have employed MD simulations of the M2 helix nanopore which corresponds to the pore lining segment of the 5-HT_3_ receptor channel ^16^. This model nanopore forms an ideal system for studying hydrophobic gating because it embodies many aspects of the wetting/de-wetting behaviour observed in complex ion channel proteins ^38^, whilst remaining sufficiently simple to allow functional dissection by an extended series of atomistically detailed simulations (Fig. 1) ^39^.

**Figure 1:**
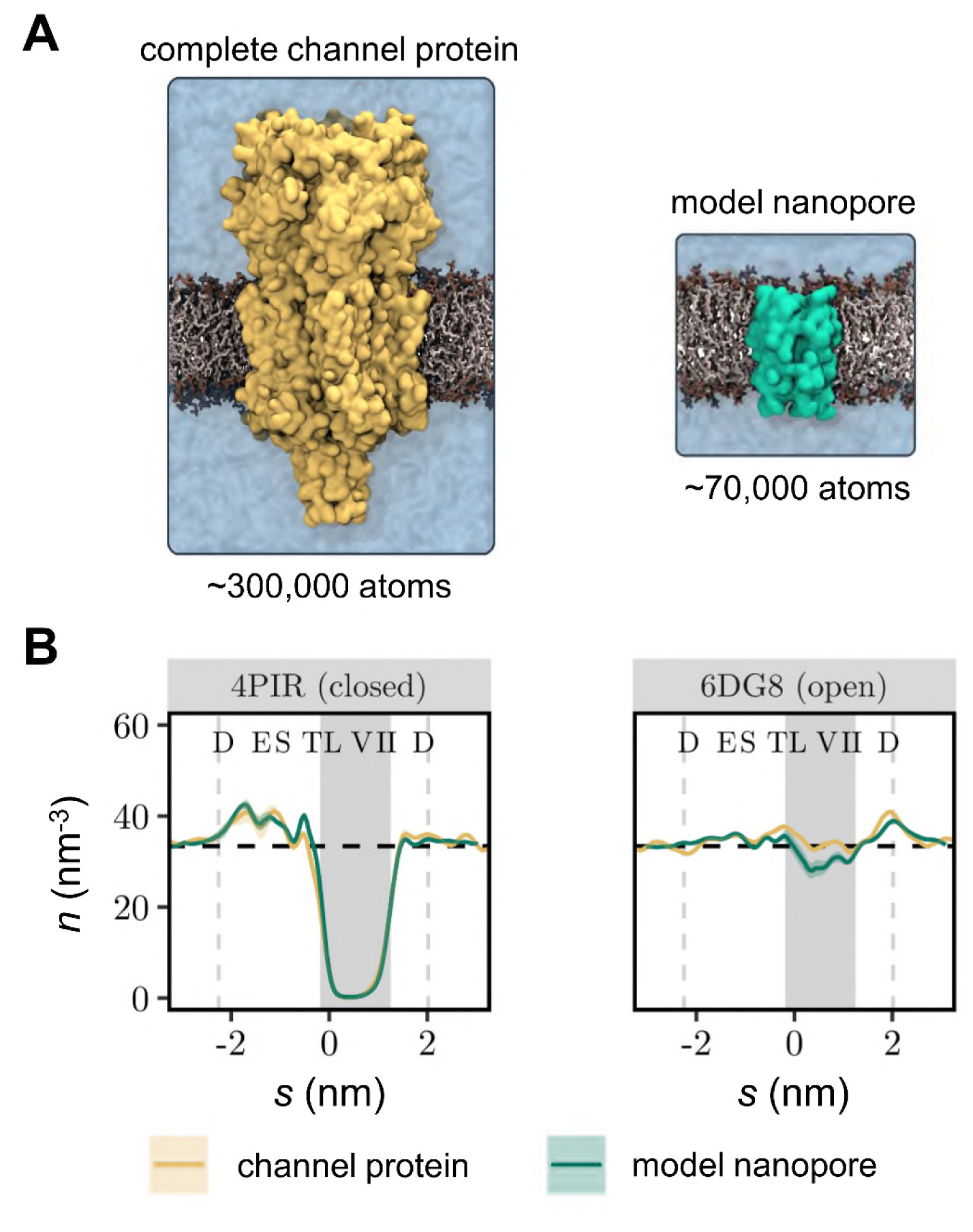
Influence of nanopore structural model on hydrophobic gating. (**A**) Simulation system based on the complete 5-HT_3_ receptor structure (~300,000 atoms for protein, bilayer and solution) compared to a much smaller simulation system (~70,000 atoms) based on the M2 helix bundle nanopore. Both structures represent the closed state (PDB ID: 4PIR) of the channel. (**B**) Time-averaged water density profiles of the complete channel protein (yellow lines) compared with the M2 helix nanopores (green line) compared for a closed state and open state of the 5-HT_3_ receptor. Vertical dashed lines indicate the extent of the M2 helix bundle pore. The shaded background represents the hydrophobic gate region, while the dashed horizontal line represents the density of bulk water (33.37 nm^−3^). Confidence bands represent the standard deviation over three independent MD simulations.

To evaluate our expectation that the model nanopore can capture the hydration state behaviour of the larger parent channel protein, we compared simulations (3 × 50 ns) of the complete 5-HT_3_ receptor in closed (PDB ID: 4PIR) ^40^ and open (PDB ID: 6DG8) ^16^ states with simulations (3 × 150 ns) of the corresponding model nanopores. In the latter simulations, only the pore-lining M2 helices of the channel were included. Both the complete protein channel and the model nanopores were embedded in a phospholipid bilayer and solvated in a ~0.15 M NaCl electrolyte using the mTIP3P water model.

These simulations show that, in the absence of an electric field, the resulting water density profiles (Fig. 1B) through the transmembrane pore regions are nearly identical for the complete channel and for the nanopore simulation systems. In both cases, the closed state (PDB ID: 4PIR) conformation that had previously exhibited significant dewetting ^38^, still favours the vapour state even when only the M2 helix nanopore is simulated. In the open conformation (PDB ID: 6DG8), the time-averaged water density in the hydrophobic gate region of the M2 helix nanopore falls slightly below the value observed for the complete channel structure, reflecting the fact that the M2 helix nanopore intermittently dewets. However, the average water density remains close to the bulk value in these simulations, suggesting the influence of the structural model on the hydration equilibrium is limited and not sufficiently strong enough to bias an otherwise hydrated channel towards the vapour state. These results therefore provide confidence these different conformations of the M2 helix nanopore can be used as a suitable model system to quantify the relationship of pore radius, electric field, and water model in the electrowetting and conductance behaviour of a biologically relevant nanopore.

### MD Simulations of the Model Nanopore in an Electric Field

We therefore examined the effect of an electric field on the liquid-vapour equilibrium that exists within the hydrophobic barrier of the M2 helix nanopore that is based on the closed state structure of the 5-HT_3_ receptor (PDB ID: 4PIR) ^40^ (Fig. 2A). Without an electric field, this nanopore remained de-wetted throughout most of a 150 ns long simulation (as seen previously ^38^) and only entered the wetted state for a small number of very brief periods (Fig. 2B). Correspondingly, the time-averaged water density profile (Fig. 2C) demonstrates that the hydrophobic gate is primarily devoid of bulk phase water and exists in a vapour state. When an electric field of ±100 mV/nm was imposed, the channel exhibited frequent liquid-vapour oscillations (Fig. 1B). In this situation, the time-averaged density of water in the hydrophobic gate adopts an intermediate value between de-wetted and hydrated channel (Fig. 1C). However, it is important to note the relationship between the units of field strength (i.e. mV/nm) used here to describe the external electric field with the values of transmembrane potential commonly measured experimentally (i.e. mV). In our simulation system, a field strength of 100 mV/nm that induces wetting in this pore (see Fig.2) corresponds to a potential difference *ΔV* of ~0.85 V across the membrane; this is calculated from *ΔV* = *EL*_*z*_, where *E* is the external field imposed along the *z* axis of a simulation box of dimension *L*_*z*_, as discussed in ^41^. Notably, this value is almost an order of magnitude greater than the typical transmembrane voltages experienced physiologically (~0.1 V).

**Figure 2:**
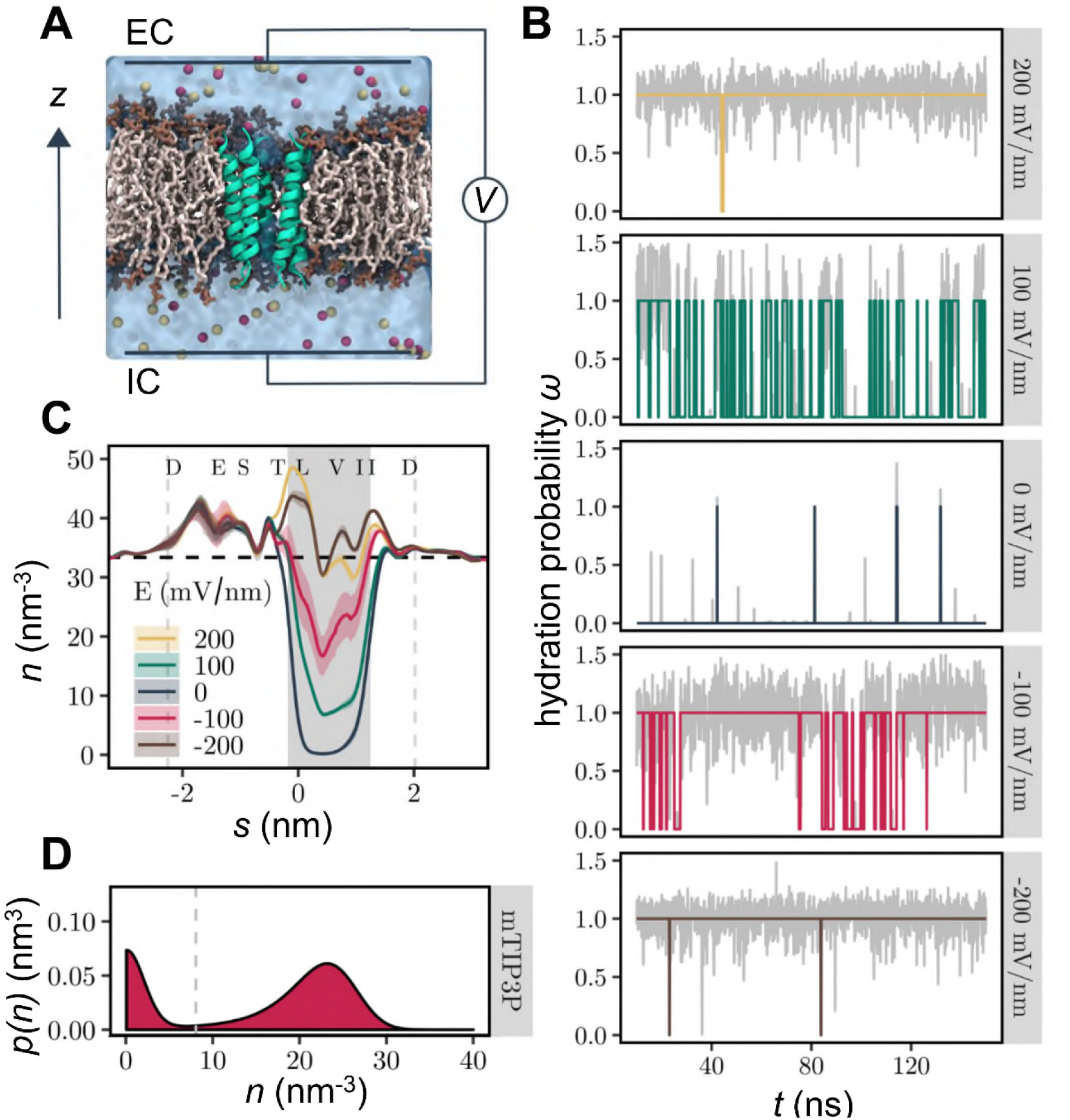
Electrowetting of the hydrophobic gate in the nanopore. **A** The M2 helix bundle nanopore (green) is embedded in a lipid bilayer (brown). Water is represented as a transparent surface and Na^+^ and Cl^−^ ions are shown as red and yellow respectively. The effect of a transmembrane potential is modelled by applying a constant electric field to all atomic charges in the system. The transmembrane voltage is reported as *V*_IC_ – *V*_EC_ where IC = intracellular (negative *z*) and EC = extracellular (positive *z*). (**B**) Time series of hydration probability (*ω*) at five different electric field strengths. The grey line in the background represents the minimum water density in the hydrophobic gate normalised to the density of bulk water. (**C**) Time-averaged water density profiles as a function of electric field strength. Confidence bands represent the standard deviation over three independent repeats. Shaded background and vertical dashed lines indicate the extent of the hydrophobic gate and transmembrane domain respectively. The horizontal dashed line represents the density of bulk water. (**D**) Kernel density estimate of water density in the hydrophobic gate region aggregated over time and electric field strength. The dashed grey line indicates the threshold density for classifying the state of the channel as closed, *ω*(*t*) = 0, or open, *ω*(*t*) = 1. Data shown are based on simulations of the M2 helix nanopore in the closed state using the mTIP3P water model.

Interestingly, we also observed that a field strength of −100 mV/nm (i.e. one in which the intracellular face of the membrane is at a negative potential relative the extracellular face; Fig. 1A) has a greater effect than a field of +100 mV/nm, indicating that the hydration probability of the pore can depend on the direction of the electric field as well as on its magnitude. At higher magnitude electric fields of ±200 mV/nm, the channel pore exists almost exclusively in the wetted state irrespective of field direction (Fig. 2B) and the average water density approaches that of bulk water (Fig. 2C).

An external electric field can therefore be used to control the phase behaviour of nanoconfined water within a biological nanopore by influencing the relative probability of the liquid and vapour states. The probability distribution of minimal water density is bimodal (Fig. 2D; SI Fig. S1), and hence the underlying free energy landscape has two minima (as discussed previously ^4, 7, 42^), with one peak near zero and a second peak near the density of bulk water. Importantly, this also legitimises our theoretical treatment of the system using a two-state model.

### Dependence of Hydration Probability on Electric Field

The response of the time averaged pore hydration probability, 〈*ω*〉, to an external electric field, *E*, was compared for the M2 bundle nanopore in conformations corresponding to closed (PDB ID: 4PIR) ^40^ and open (PDB ID: 6DG8) ^16^ states of the 5-HT_3_ receptor. The hydration behaviour of the nanopore was also compared for four different water models (Fig. 3). For the closed state, the hydration probability vs. *E* response curve is sigmoidal: the hydration probability is near zero in the absence of a field, increasing to a value of one at fields of high magnitude. The threshold for hydration also varies. For example, with the mTIP3P water model pore wetting is observed at a field strength which is ∼50 mV/nm weaker than with any of the other models. However, once pore wetting begins, the transitional regime of intermediate hydration occurs within a further increase of ∼ 100mV/nm for all four water models. In case of the open state structures, the response of hydration probability still resembles a (double) sigmoid, but hydration probability does not start at zero, because the pore is already partially hydrated even in the absence of an electric field.

**Figure 3:**
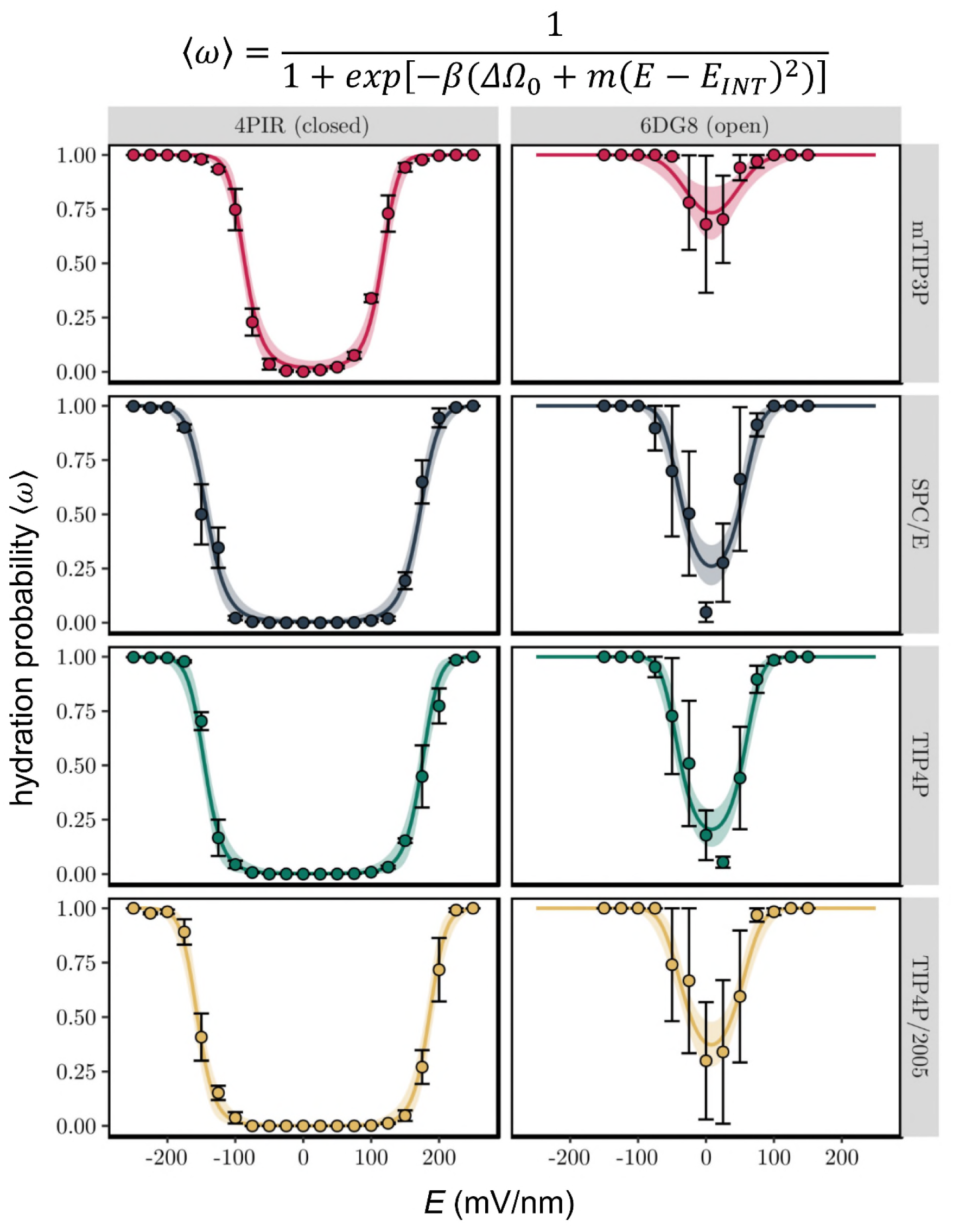
Hydration probability 〈*ω*〉 as a function of the external electric field, *E*. Discrete data points and error bars represent the mean hydration probability and its standard error over three independent simulations respectively. Solid lines are the result of a Bayesian nonlinear multilevel model (equation given above the figure and in the main text) fitted to the data (see Table 1 and SI Fig. S3 for fitted parameters). The shaded area indicates the 95% confidence interval of the fit. Simulations were performed on the M2 helix nanopore from the closed (PDB ID: 4PIR) and open (PDB ID: 6DG8) conformations of the 5-HT_3_ receptor and employed four different water models.

It is important to point out that in the transitional regime the rate of increase in hydration probability appears similar for both positive and negative electric fields, irrespective of the water model. However, the overall curve is shifted towards the right, which makes it asymmetric with respect to *E* = 0. This is most clearly seen for the closed state, where the threshold voltage for the onset of pore hydration is ∼25 mV/nm higher at positive fields than for negative fields. Due to the smaller number of data points in the de-wetted regime, this effect is not as clear for the open state although, consistent with this effect, the minimum point of the pore hydration probability curve appears shifted to slightly positive fields.

**Table 1:** Bayesian estimates of parameters for a model of pore hydration probability as a function of *E*-field.

| PDB ID | water model | $\Delta\Omega_o$ (k<sub>B</sub>T) | $m$ (k<sub>B</sub>T nm<sup>2</sup>V<sup>-2</sup>) | $E_{\text{INT}}$ (mV nm<sup>-1</sup>) |
| --- | --- | --- | --- | --- |
| 4PIR | mTIP3P | $-4.03 \pm 0.68$ | $396 \pm 67$ | $14.8 \pm 1.4$ |
| 4PIR | SPC/E | $-5.51 \pm 0.88$ | $225 \pm 36$ | |
| 4PIR | TIP4P | $-6.47 \pm 1.32$ | $253 \pm 52$ | |
| 4PIR | TIP4P/2005 | $-7.08 \pm 1.31$ | $243 \pm 45$ | |
| 6DG8 | mTIP3P | $1.03 \pm 0.34$ | $436 \pm 171$ | $7.9 \pm 2.0$ |
| 6DG8 | SPC/E | $-1.06 \pm 0.25$ | $668 \pm 139$ | |
| 6DG8 | TIP4P | $-1.37 \pm 0.28$ | $695 \pm 126$ | |
| 6DG8 | TIP4P/2005 | $-0.53 \pm 0.23$ | $571 \pm 122$ | |

The probability of hydration of a nanopore in the absence of an *E*-field can be described by:

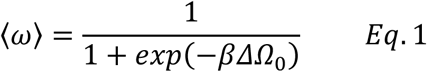

where 〈*ω*〉 = the time-averaged hydration probability, *ΔΩ*_0_ = *Ω*_*V*_ − *Ω*_*L*_ *i.e.* the difference between the free energies of the liquid and vapour states in the absence of an *E*-field, and *β* = *1/k*_*B*_*T* ^7, 43^. When *ΔΩ*_*0*_ < 0, the pore is hydrophobic and favours a vapour state, but when *ΔΩ*_*0*_ > 0, the liquid (*i.e.* hydrated) state is favoured. To include the effect of the electric field, *E*, the free energy term ^28^ can be modified giving:

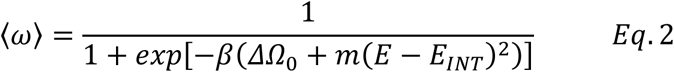

where *m* denotes the strength of the coupling between hydration probability and the magnitude of the field and *E*_*INT*_ accounts for the horizontal offset of the response curve in terms of an intrinsic electric field arising from the nanopore structure (see below). The value of *m* represents the difference in ability to store electrical energy in a water-filled high-permittivity space vs an empty low-permittivity space, and this is related *inter alia* to the wettable volume of the pore as well as the local dielectric constant of the water model. As can be seen from Fig. 2, this model is in good agreement with the simulation data.

Model parameters are given Table 1 (with errors estimated from the Bayesian posterior distributions shown in SI Fig. S3). These parameters enable a quantitative analysis of the effects of E-field strength, nanopore conformation, and water model on the hydration probability of the channel. Thus, the zero-field free energy difference is close to zero across all water models for the open state structure with the mTIP3P water model slightly favouring the liquid state by ~1 k_B_T, whilst the other three models have an approximately −1 k_B_T preference for the vapour state. In marked contrast, all water models strongly favour the vapour state in the closed conformation, with zero-field free energy differences ranging from −4 to −6 k_B_T. Notably, values using the mTIP3P model appear closest to the liquid state for both conformations. The strength of the coupling, *m*, between the hydration state of the pore and the electric field is also dependent on the conformation of the receptor. Apart from the mTIP3P water model, this parameter is more than twice as large for the open state than for the closed state. This difference may be attributed to the increased pore radius of the open conformation, which corresponds to a larger volume of water within the pore. There appears to be no clear trend for how *m* varies across water models. However, it should be noted that this parameter is sensitive to intermediate values of openness (〈*ω*〉 ≈ 0.5) and therefore may be affected by the larger error bars found in the transitional regime.

The horizontal offset to more positive potentials of the hydration probability is quantified by *E*_*INT*_. In the closed conformation, *E*_*INT*_ is 14.8 ± 1.4 mV/nm. This reduces to 8.0 ± 2.0 mV/nm in the open conformation. Due to the sign convention this intrinsic field points from the extracellular to the intracellular domain. It therefore adds to the effective magnitude of an inward pointing external field (*E < 0*), while effectively reducing the impact of an outward pointing external field (*E > 0*). In this thermodynamic model, the shifting parameter, *E*_*INT*_, is associated with the electric field arising from the charged amino acids of the channel protein, i.e. the intrinsic electric field of the 5-HT_3_ receptor M2 bundle. This was demonstrated by determining the electrostatic potential of the M2 bundle of the closed conformation by numerically solving the Poisson-Boltzmann equation (SI Fig. S2A). The overall shape of the resultant electrostatic potential profile indicates that there is a substantial potential gradient between the intra- and extracellular openings of the M2 helix bundle, from −160 mV in the intracellular mouth well (IC in SI Fig. S2) to −100 mV in the extracellular well (EC in SI Fig. S2, when a lipid bilayer is included in the calculation). This corresponds to an electric field of 20 mV/nm, which is close to the *E*_*INT*_ value of ~ 15 mV/nm found from the fit (Table 1). The relative distribution of charged amino acids (SI Fig. S2B) therefore provides a molecular explanation of the resultant electrostatic profile.

### Ion conduction through the nanopore

In addition to determining the influence of the electric field on pore hydration, we also measured the resultant ion conduction through the channel (Fig. 4; also SI Fig. S4). For the closed (i.e. dehydrated) state the current due to either ion species (i.e. Na^+^ or Cl^−^) is approximately zero at low electric fields and begins to increase in magnitude only when the field exceeds the critical value required to hydrate the hydrophobic gate. Following this initial nonlinearity, the current then increases linearly with field strength for large electric fields. This implies that once the pore is hydrated, it acts as an Ohmic resistance. For the open state, the conductance is approximately Ohmic at lower field strengths (> 50 mV/nm) reflecting the greater hydration of the pore. Interestingly, if we calculate the slope conductance at the highest field strengths (i.e. *G_max_ = dI/dV* at *E* = +200 mV/nm) for both the closed and open states, these values of *G*_*max*_ are of comparable magnitude (except perhaps for TIP4P/2005). This suggests that once this model nanopore is fully hydrated and at relatively high *E*-fields, the overall shape and size of the pore dominates the limiting conductance, as discussed in early theoretical treatments of ion channel conductance ^44^. Importantly, these results also indicate that pore de-wetting prevents the formation of an ionic current in the closed state. This in turn implies that the permeation of ions itself does not cause pore hydration, given that the threshold field strength for hydration is dependent on the water model. This is in agreement with previous simulations of model nanopores, where hydration was seen to occur a fraction of a nanosecond before ion permeation once a transbilayer *E*-field was imposed ^35^.

**Figure 4:**
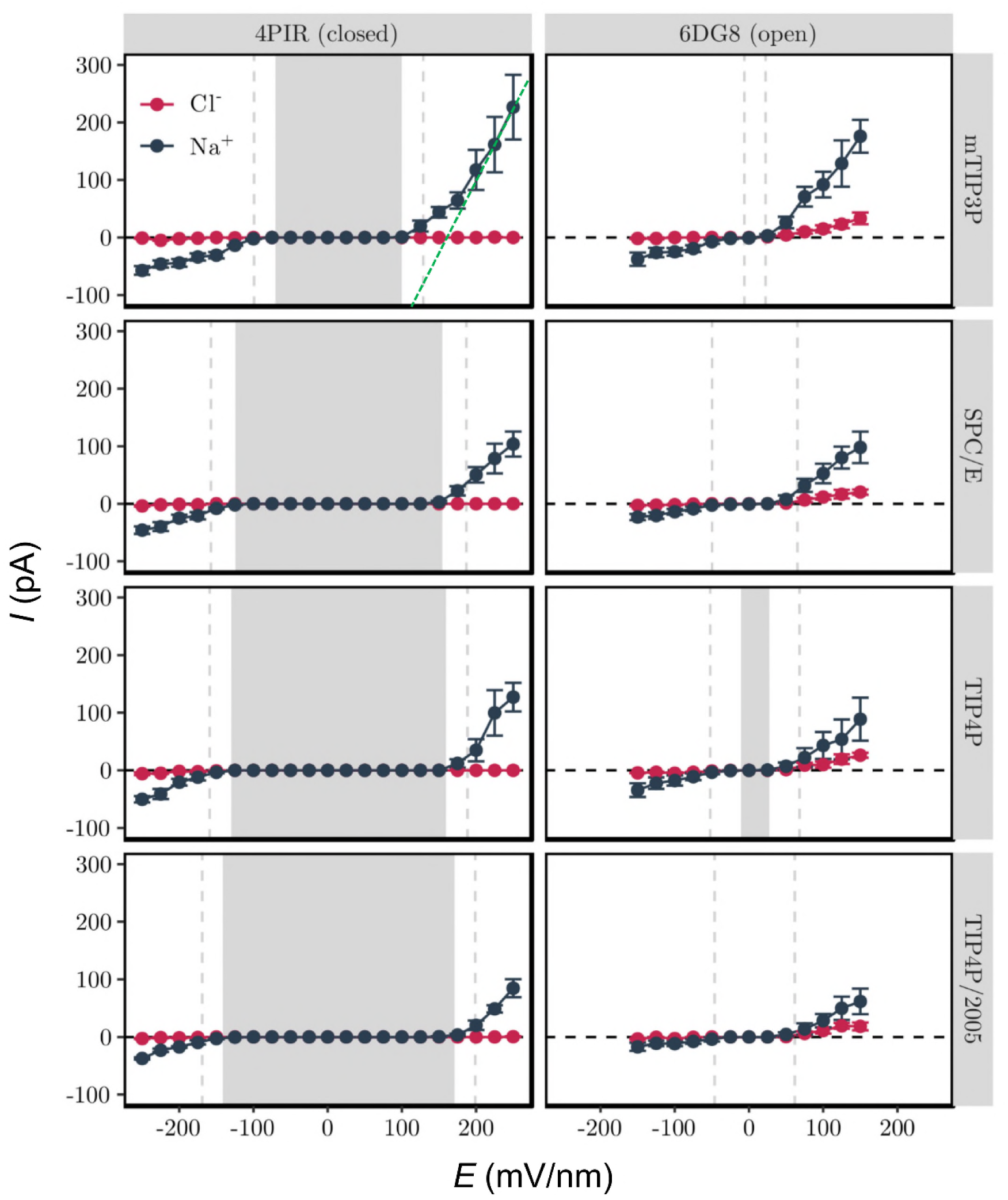
Ionic current through the 5-HT_3_ receptor M2 helix nanopore in dependence of the external electric field. Background shading indicates the regime where the hydrophobic gate is mostly dewetted (〈*ω*〉 < 0.25), while the vertical dashed lines indicate the transition to a mostly hydrated (〈*ω*〉 > 0.75) pore. The green dotted lines shown for the mTIP3P closed state plot represent the limiting Ohmic conductance (*G*_*max*_) at high *E*-field (see text for further details).

The value of *G*_*max*_ from our simulations exceeds ~100 pS (Fig. 4) which is is more than two orders of magnitude greater than the physiologically observed single channel conductance (~1 pS) ^45^. However, this difference likely reflects both the high *E-*field strength used in these simulations, and also the absence of the intracellular and extracellular domains of the protein, which reduce the conductance substantially. Nevertheless, the Na^+^ current is significantly larger than the Cl^−^ current, in agreement with the cation selectivity of the 5-HT_3_ receptor ^46^. This is most likely due to the negatively charged residues near the intracellular (D −4′, E −1′) and extracellular (D20′) openings of the M2 helix bundle (SI Figure S2B), which make entry of Cl^−^ ions into the pore highly unfavourable, whilst remaining attractive for Na^+^.

### Kinetics of Liquid-Vapour Transitions

Having shown that an external E-field influences the hydration probability of a nanopore by changing the free energy difference between its liquid and vapour states, we next explored how the free energy difference is controlled by the underlying rate constants. The kinetics of liquid-vapour oscillations inside a hydrophobic gate can be examined in terms of the mean survival times of the liquid state, τ_l_, and vapour state, τ_v_, as they vary with the free energy difference between these two states. In Fig. 5. it can be seen that τ_l_ grows exponentially with ∆Ω, while τ_v_ decreases exponentially at approximately the same rate. This trend is seen for both the closed and open states of the nanopore channel and across all four water models.

**Figure 5:**
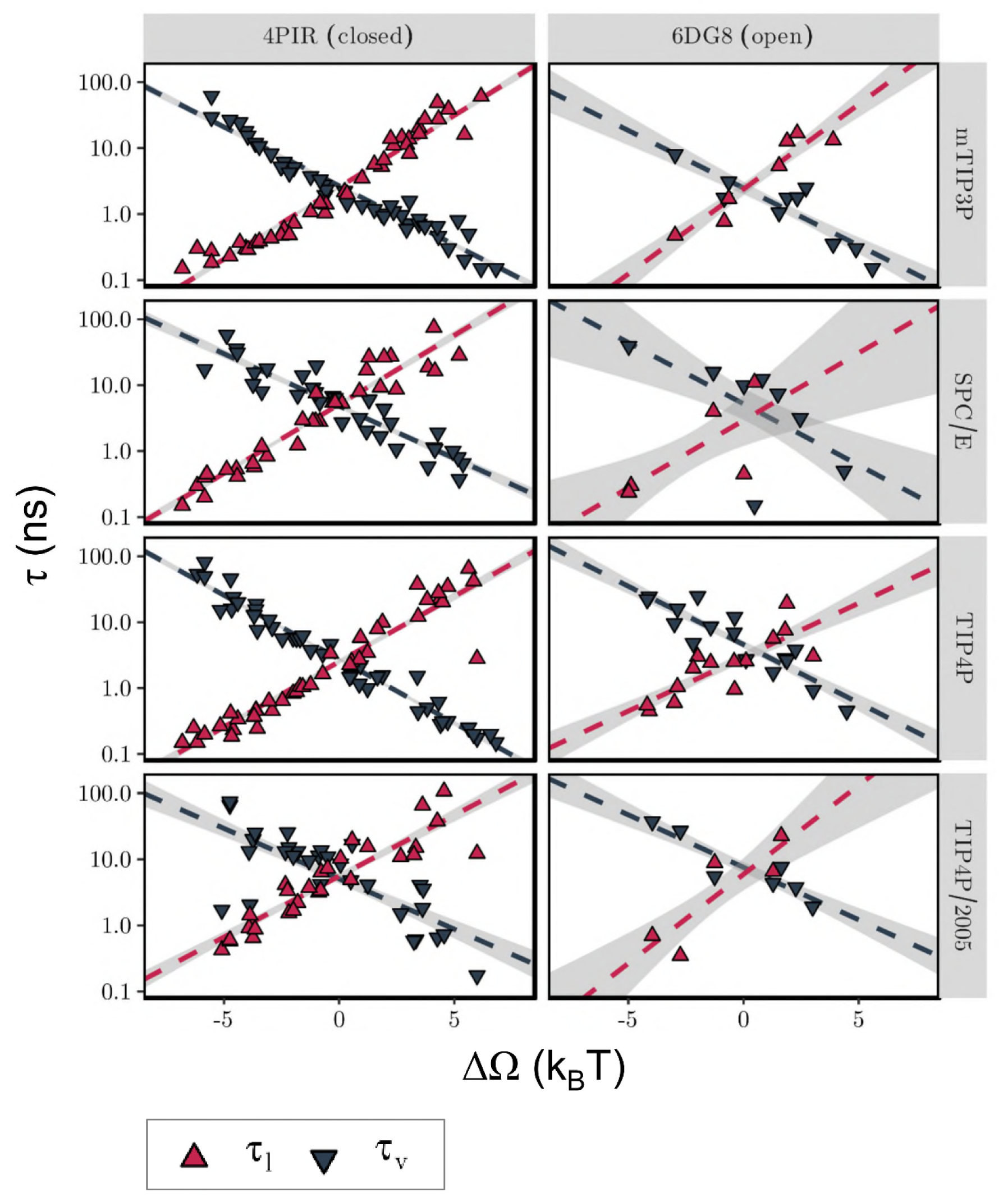
Kinetics of liquid-vapour transitions under the influence of an external electric field. The dependence of the mean survival times of the liquid (τ_*l*_ red) and vapour (τ_*v*_ black) states on the free energy difference (∆Ω) between these states is shown. Each data point corresponds to an individual simulation, while the dashed line represents a linear fit to all data points, with confidence bands indicated as shaded background. The free energy difference was estimated from the mean hydration probability, while the survival times were determined from the time series of hydration probability. Simulations with fewer than two observed transitions are not included in this plot.

The lines in Fig. 5 necessarily cross one another at ∆Ω = 0, as an equal probability for both states implies equal survival times. The crossover point thus defines a characteristic time scale for liquid-vapour oscillations, which for the M2 helix nanopore has a value of τ_lv_ = τ_l_ (∆Ω = 0) = τ_v_ (∆Ω = 0) ≈ 5 ns. Importantly, this value is nearly identical for both the closed and open states of the nanopore, despite their different pore radii. Furthermore, ∆Ω = 0 occurs at very different magnitudes of the electric field. This indicates that a time scale of ~5 ns is likely to characterise liquid-vapour oscillations of all pores that share the characteristic dimensions and surface hydrophobicity of the M2 helix nanopore considered here. Such slow behaviour (water has a typical H-bond lifetime of the order of psec) indicates these oscillations arise collectively, reminiscent of e.g. ultraslow reorientation of water wires with nanotubes ^47^ or in extended networks at the surface of a protein ^48^ There is some slight variation of this oscillation time scale τ_lv_ across water models, but it always falls in the range between 1 ns and 10 ns, indicating a limited force field sensitivity.

This also suggests that a total simulation time which is an order of magnitude larger than the characteristic time scale (i.e. 10 τ_lv_ ≈ 50 ns or above) is sufficient to characterise the hydration equilibrium behaviour of the M2 helix nanopore systems. It is also worthwhile to compare the duration of the liquid and vapour states with the typical dwell time of ions inside the channel pore (SI Fig. S5). For ions that pass the closed conformation in its partially hydrated state (〈*ω*〉 = 0.25 to 0.75), the average dwell time is comparable to the typical time scale of liquid-vapour oscillations (τ_lv_ ≈ 5ns). This implies that the intermittent periods during which the channel exists in the liquid state last sufficiently long enough to allow the passage of individual ions. For the open state, the typical ion dwell time lies below ∼1 ns. This is less than the average lifetime of the liquid state, which exceeds ~1 ns. Consequently, even though most water models predict some degree of dehydration of the open state nanopore in the absence of an electric field (Fig. 3), this state is still expected to allow the regular passage of ions, and so is in agreement with our previous annotation of this structure as a functionally open state ^38^.

## Conclusions

In summary, we have shown that a transbilayer electric field can induce wetting of the hydrophobic gate of a model nanopore derived from the 5-HT_3_ receptor. However, a field strength of ∼100 mV nm^−1^ is required to start to hydrate the pore of the closed conformation of the channel. This corresponds to a potential difference of ∼0.9 V, i.e. substantially greater than the transmembrane voltage an ion channel would normally experience under physiological conditions. We also present a simple model which can quantitatively describe the observed increase in pore hydration probability with increasing electric field, with the asymmetry of the hydration probability response curve accounted for by an offset parameter, *E*_*INT*_, linked to the intrinsic electric field generated by the unequal distribution of charged amino acid residues along the length of the pore.

It is also useful to reflect on possible extensions to the current studies. One possibility would to be extend the comparison of water models to include polarizable models ^39^. However, as yet for e.g. AMOEBA ^49–50^ external E-fields are not available in the GPU-accelerated OpenMM toolkit ^51–52^, because the external-field-induced-dipole interaction is not implemented. Furthermore, it is likely that anisotropic atomic polarizability of the water molecule ^53^ may be needed for more accurate treatment of electrowetting. Another possible extension to this work would be to explore electrowetting in a complete protein channel. This would be computationally demanding, but would be of interest e.g. in terms of how a more complex distribution of charge within a protein might influence *E*_*INT*_.

The finding that the threshold voltage required for hydrating a hydrophobic gate depends on the orientation of the electric field also provides an attractive perspective for the design of artificial nanopores. Our results suggest that an excess of only five elementary charges (from amino acid sidechains) near one opening of the pore is sufficient to create a substantial intrinsic electric field acting in the transmembrane direction. Therefore, by functionalising the openings of a hydrophobic pore with static charges ^54^, it should be possible to design a nanopore that permits ion transport in only one direction, thus rectifying ionic currents. Importantly, this rectifying property would not only apply to ions, but also to water molecules. This not only extends the concept of the electric-field induced smart water gates ^55^, but may also enable the creation of membrane systems that are semi-permeable to water in a switchable fashion.

## Methods

### Molecular Dynamics Simulations

MD simulations employing a constant electric field were performed in GROMACS 2018 ^56^ using the protocols described below. Protein structures were obtained from the PDB and missing atoms were added using WHAT IF ^57^. A 100 ns long MD simulation using the MARTINI force field ^58^ of the M2 helix bundle nanopore together with DOPC lipids and water was used to place the nanopore in a lipid bilayer ^59^. During this coarse-grained simulation lipid molecules self-assemble into a bilayer structure around the protein. Following the self-assembly simulation, the protein-bilayer system was converted back to an atomistic representation and was solvated in a 0.15 M NaCl solution. The system was then equilibrated through a 10 ns long MD simulation employing the mTIP3P water model ^60^ and the CHARMM36m force field ^61–62^.

### Constant Electric Field Simulations

Production simulations were carried out using the CHARMM36m protein force field ^61^ and associated lipid parameters ^61–62^ in conjunction with the mTIP3P ^60^, SPC/E ^63^, TIP4P ^60^ or TIP4P/2005 ^64^ water models. During these simulations, the effect of an external electric field, *E*, was modelled through an additional force, *q*_*i*_*E*, acting on each atomic partial charge, *q*_*i*_. The electric field was applied in the *z*-direction perpendicular to the plane of the membrane. Figure 2A illustrates this simulation setup.

It has been shown ^41^ that under periodic boundary conditions the application of an external field as described above is equivalent to the application of a transmembrane voltage, such that the voltage drop across the membrane is given by *V = EL_z_*, where *L*_*z*_ is the length of the unit cell in the z-direction. For the system considered here *L*_*z*_ ≈ 8.5 nm.

The magnitude of the electric field, *E*, was varied in steps of 25 mV nm^−1^ between −250 mV nm^−1^ and 250 mV nm^−1^ for the closed state structure (PDB ID: 4PIR) and between −150 mV nm^−1^ and 150 mV nm^−1^ for the open state structure (PDB ID: 6DG8). In terms of transmembrane potential, this corresponds to a step size of 0.2125 V and maximum transmembrane voltages of ± 2.125 V and ± 1.275 V respectively. In each case, the range was chosen to ensure that a complete transition from a (partially) de-wetted to a fully hydrated pore could be observed.

Preliminary simulations were carried out to assess the stability of the simulation system under electric fields of this magnitude. In order to assess pore hydration properties under high-field (|E| > 150 mV nm^−1^) conditions (under which conditions electroporation may occur ^65^), a harmonic restraining force of 1000 kJ mol^−1^ nm^−2^ was placed on the heavy atoms of all lipid molecules (after the initial 10 ns long equilibration period), which prevented electroporation.

### Simulation Details

Three independent simulations of duration 150 ns for the closed conformation (PDB ID: 4PIR) and 50 ns for the open conformation (PDB ID: 6DG8) were carried out at each value of the electric field. Time integration was performed using a leapfrog integrator with a step size of 2 fs. Covalent bonds to hydrogen atoms were constrained using the LINCS algorithm ^66^ and water model geometry was enforced using the SETTLE method ^67^. The conformation of the protein was maintained close to its experimentally determined structure through the application of a harmonic restraining potential of 1000 kJ mol^−1^ nm^−2^ to all Cα atoms.

Long-range electrostatic interactions were treated through the smooth PME method ^68^ employing a real space cutoff of 1.0 nm and a Fourier spacing of 0.12 nm, with charge interpolation onto the grid through fourth order B-splines. Lennard-Jones interactions were switched off between 1.0 nm and 1.2 nm and a long-range dispersion correction was applied to energy and pressure. All simulations were carried out in the isothermal isobaric (NPT) ensemble. Temperature was maintained at 310 K through a velocity rescaling thermostat ^69^ with a coupling constant of 0.1 ps, while a pressure of 1.0 bar was enforced through a semi-isotropic Parinello-Rahman barostat ^70^ with a coupling constant of 1.0 ps and a compressibility of 4.5 × 10^−5^ bar^−1^.

### Analysis

Unless otherwise specified, MD trajectories were post-processed using MDAnalysis version 0.19.0 ^71^ while further data analysis and visualisation were performed through custom scripts written in the R language (https://www.r-project.org/). Molecular visualisations were created in VMD ^72^.

As discussed above, the electrowetting of hydrophobic gates can be understood by considering the influence of an external electric field on the free energy difference between the liquid and vapour states of the pore (Eq. 2 above). Consequently, the hydration probability can be written as

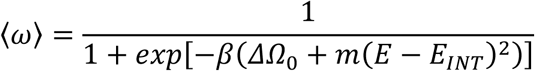

which constitutes a nonlinear model with three free parameters, namely ∆Ω_*0*_ (the free energy difference in the absence of an external electric field), *m* (which represents how strongly the probability of hydration is coupled to the electric field), and *E*_*INT*_ (the horizontal offset accounting for the intrinsic electric field due to the protein; see above for more detailed discussion).

A fit of this equation to the observed hydration probability was obtained within the framework of Bayesian multilevel modelling ^73^. Both ∆Ω_*0*_ and *m* were treated as dependent on both channel structure and water model, while *E*_*INT*_ was considered to only depend on the channel structure. The zero-field free energy difference, ∆Ω_*0*_, may take both negative and positive values and a Gaussian prior with mean *µ* = 0 and standard deviation *σ* = 10 k_B_T was used. In contrast to this, the coupling strength, *m*, is positive definite by definition. A gamma prior with shape *α* = 2 and rate *β* = 0.01 was chosen to enforce this constraint and sampling for this parameter was limited to positive values. While the intrinsic electric field, *E*_*INT*_, can in principle be both positive and negative, allowing this parameter to vary over the entire real range caused divergent transitions in the sampling process. Sampling for this parameter was therefore restricted to positive values (in accordance with the observed data) and a gamma prior with shape *α* = 2 and rate *β* = 100 was chosen.

The model was implemented in the probabilistic programming language Stan ^74^ through the brms ^75^ library in R. Posterior samples were generated through a Hamiltonian Monte Carlo ^76–77^ procedure using the No-U-Turn (NUTS) sampler ^78^. For each of ten independent Markov Chains, 10000 warm-up and 10000 sampling iterations were carried out. The NUTS sampler used a target average acceptance ratio of 0.9 and a maximum tree depth of 15.

## Acknowledgments

This work was supported by the BBSRC, EPSRC, HECBioSim, and the Wellcome Trust. Our thanks to Shanlin Rao and Charlotte Lynch for their interest in and helpful comments on this work.

## Supporting Information

Including: SI Methodological Details, and SI Figures (S1 to S5).

