## Supporting Information for "Electric Field Induced Wetting of a Hydrophobic Gate in a Model Nanopore Based on the 5-HT_3_ Receptor Channel"

### SI Methodological Details

*Poisson-Boltzmann Electrostatics*

In addition to the MD simulations described in the previous section, the electrostatic potential, φ(r), around the nanopore model derived from the 5-HT$_3$ receptor was calculated by numerically solving the non-linear Poisson-Boltzmann equation using the Adaptive Poisson-Boltzmann Solver (APBS) [1]. Poisson-Boltzmann calculations were carried out on the M2 helix bundle of the closed conformation of the channel (PDB ID: 4PIR), both in the presence and absence of a DOPC lipid bilayer. A multigrid finite differencing method was employed: the coarse grid was 2.5 times as large as the bounding box around the M2 helix bundle and had a grid spacing of 0.17 nm, while the fine grid was 1.5 times as large as the bounding box and had a grid spacing of 0.10 nm.

For consistency with the MD simulations charges were assigned to the atomic sites according to the CHARMM36m force field and were discretised onto the grid through cubic B-spline interpolation. An electric permittivity of 2 for the protein and 78.5 for the solvent were used and calculations were carried out at a temperature of 310 K. A 150 mM NaCl electrolyte was used as solvent and alongside ionic Born radii of 0.1680 nm for $Na^+$ and 0.1937 nm for $Cl^-$. A one-dimensional electrostatic potential profile, φ(s), was calculated from the three dimensional potential field, φ(r), resulting from the Poisson-Boltzmann calculation, along the centre line spline curve, **s**(s), which represents the likely path along which an ion would permeate the channel [2].

*Hydration Probability and Kinetic Constants*

CHAP [2] was used to calculate a water density profile, *n(s,t),* along the channel pore and the minimal instantaneous water density and a threshold crossing method was used to determine a time series of channel openness [3]. The time-averaged openness was subsequently calculated as a measure for the probability of finding the channel pore in a hydrated state $\langle\omega\rangle$. A thermodynamic model predicting the dependence of $\langle\omega\rangle$ on the external electric field, *E*, is described in the main body of the text. From $\langle\omega\rangle$, the free energy difference between the liquid (*l*) and vapour (*v*) states was estimated as

$$\Delta\Omega = \Omega_v - \Omega_l = -k_B T \ln\left(\frac{1-\langle\omega\rangle}{\langle\omega\rangle}\right)$$

where $k_B$ is the Boltzmann constant and *T* is temperature.

In addition, the mean survival times of the liquid state, $\tau_l$, and vapour state, $\tau_v$, were determined. These two constants are related to the probability of hydration through

$$\frac{1-\langle\omega\rangle}{\langle\omega\rangle} = \frac{\tau_v}{\tau_l}$$

Survival times were estimated directly from the time series of $\omega(t)$ as the average duration of the liquid ($\omega(t)$ = 1) and vapour states ($\omega(t)$ = 0) respectively. The initial and final periods of the time series were excluded from this calculation, as it is unclear for how long beyond the end of simulation a state would have persisted (or for how long before the beginning of a simulation it can be assumed to have persisted). Note that this implies that at least two liquid-vapour transitions need to be observed during a trajectory in order to estimate survival times. Furthermore, no time scales longer than the duration of a simulation (150 ns for simulations of the closed and 50 ns for the open state conformation of the nanopore) or shorter than the sampling time (0.1 ns for all simulations) could be estimated.

*Ion Transport and Ionic Currents*

Ion transport was analysed through custom scripts based on the MDAnalysis library [4]. For each value of the electric field, $E$, the net current, $I$, through the ion channel pore was calculated. The simulation cell was divided into three distinct domains representing the intra- and extracellular spaces as well as the pore. If an ion was located in a cylindrical region centred on the protein, it was considered to be inside the pore and was assigned the domain index $\gamma = 0$. An ion located outside this domain was assigned to the extracellular domain ($\gamma = +1$), if its z-coordinate placed it above the middle of the lipid bilayer, and to the intracellular domain ($\gamma = -1$) otherwise.

Over the course of a simulation, an ion may jump between these three domains and conduction events can be identified from the sequence of domain indices. In order to move directly between the extra- and intracellular domains ($\gamma = +1 \rightarrow \gamma = -1$ or $\gamma = -1 \rightarrow \gamma = +1$), an ion would have to either cross the lipid bilayer or transition through the periodic boundary. These events were disregarded and only true pore crossing events ($-1 \rightarrow 0 \rightarrow +1$ or $+1 \rightarrow 0 \rightarrow -1$) were counted towards the amount of charge conducted by the ion channel. Each such conduction event was assigned a crossing number, $\eta$, based on whether the particle moved from the intra- to the extracellular domain (i.e. in positive z-direction, $\eta = +1$) or vice versa ($\eta = -1$). If an ion entered the pore ($\gamma = \pm 1 \rightarrow \gamma = 0$) at time $t^{in}$ and left it ($\gamma = 0 \rightarrow \gamma = \pm 1$) at time $t^{out}$, the time at which a crossing event occurred was taken to be $t^{cross} = (t^{out} + t^{in})/2$. Correspondingly, the ion dwell time inside the pore is given by $T_{pore} = t^{out} - t^{in}$ .

Each ion may cross the pore multiple times and thus $t^{cross}{}_{i,j}$ denotes the time at which the $i$-th ion crossed the pore for the $j$-th time, while $M_i$ denotes the overall number of crossings of ion i. The cumulative amount of charge transported through the channel pore between the beginning of the simulation and time t was then calculated according to

$$Q(t) = \sum_{i=1}^{N} \sum_{j=1}^{M_i} \int_0^t q_i \eta_{i,j} \delta\left(t' - t_{i,j}^{cross}\right) dt'$$

where $q_i$ is the charge of the $i$-th ion, $N$ is the total number of ions, and $\delta(t)$ is the Dirac delta function. The rate of ion transport was approximately constant and thus $Q(t)$ grows linearly over time. A linear model of the form $Q(t) = It$ was therefore fit to the data using R (https://www.r-project.org/) where the proportionality constant $I$ represents the electric current through the channel pore. The individual contribution of $Na^+$ and $Cl^-$ ions to the net current was evaluated by carrying out the outer sum over only a single ion species.

*Simulation and data analysis scripts*

These can be found at https://github.com/XXXXX.

*References*

1. Baker, N. A.; Sept, D.; Joseph, S.; Holst, M. J.; McCammon, J. A., Electrostatics of nanosystems: application to microtubules and the ribosome. *Proc. Nat. Acad. Sci. USA* **2001,** *98,* 10037-10041.

2. Klesse, G.; Rao, S.; Sansom, M. S. P.; Tucker, S. J., CHAP: a versatile tool for the structural and functional annotation of ion channel pores. *J. Molec. Biol.* **2019,** *431*, 3353-3365.

3. Beckstein, O.; Sansom, M. S. P., Liquid–vapor oscillations of water in hydrophobic nanopores. *Proc. Nat. Acad. Sci. USA* **2003,** *100*, 7063-7068.

4. Gowers, R. J.; Linke, M.; Barnoud, J.; Reddy, T. J. E.; Melo, M. N.; Seyler, S. L.; Dotson, D. L.; Domanski, J.; Buchoux, S.; Kenney, I. M.; Beckstein, O. In *MDAnalysis: A Python package for the rapid analysis of molecular dynamics simulations,* Proceedings of the 15th Python in Science conference (SciPy 2016), Austin, TX, Benthall, S.; Rostrup, S., Eds. Austin, TX, 2016; pp 102-109.

**SI Figures**

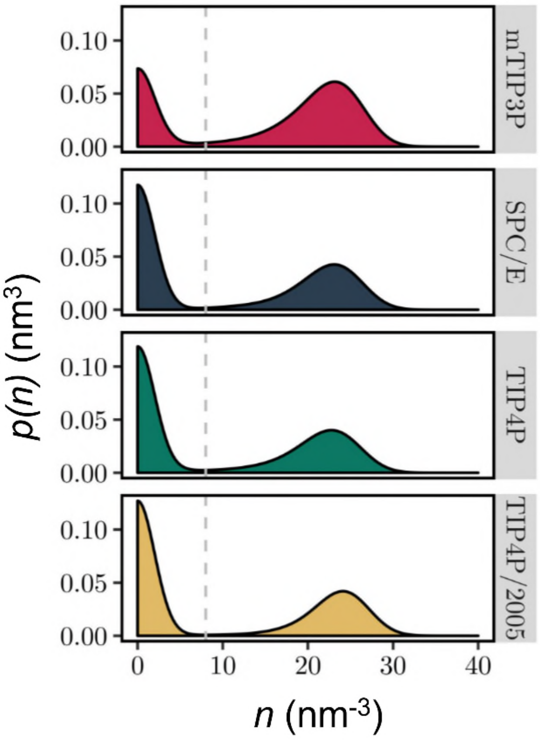


*Figure S1:*

Electrowetting of the hydrophobic gate in the 5-HT3 receptor comparing four water models. The kernel density estimate of water density in the hydrophobic gate region aggregated over time and electric field strength. The dashed grey line indicates the threshold density for classifying the state of the channel as closed ($\omega(t) = 0$) or open ($\omega(t) = 1$). Data shown are based on simulations of the M2 helix bundle of the closed conformation of the 5-HT3 receptor (PDB ID: 4PIR).

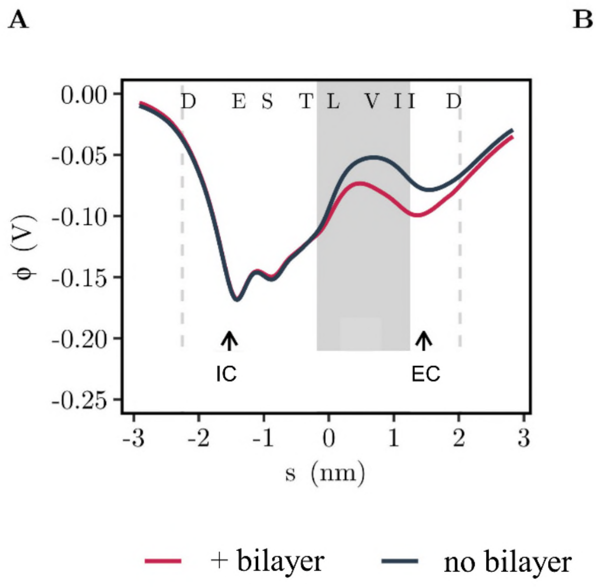


*Figure S2:*

Electrostatic potential of the M2 helix bundle of the closed state 5-$HT_3$ receptor (PDB ID: 4PIR). (**A**) Profile of electrostatic potential along the pore centre line, calculated for a helix bundle in the absence and the presence of a surrounding DOPC bilayer. (**B**) Structure of M2 helix bundle. Positively and negatively charged amino acids are shown in blue and red respectively, all other residues are coloured grey. For visual clarity, only two subunits are shown.

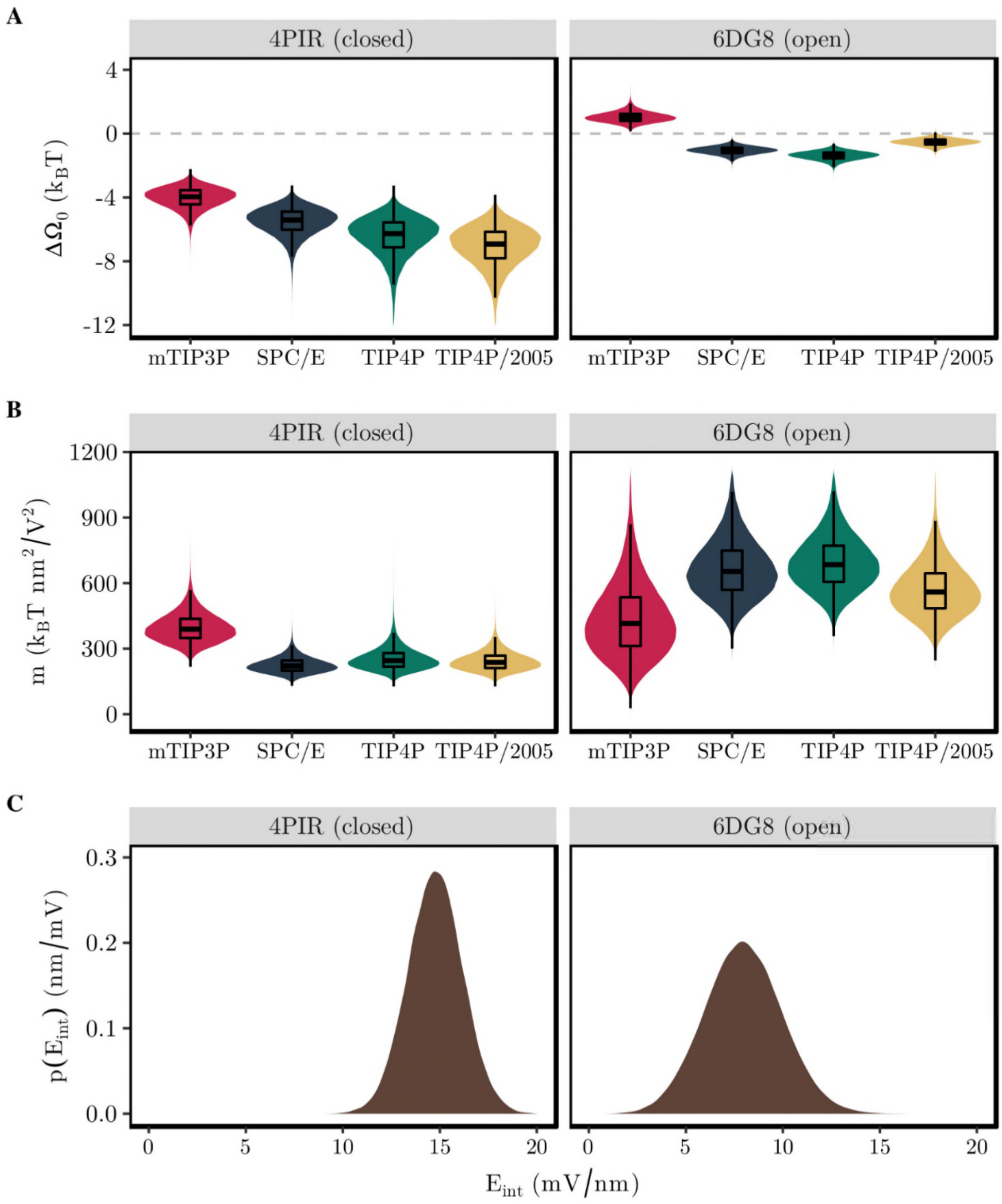


*Figure S3:*

Posterior probability of Bayesian model parameters. (**A**) Posterior distribution of zero-field free energy difference, $\Delta\Omega_0$. Data are shown as box plot superimposed on a violin plot for each combination of water model and conformational state. The dashed line indicates equal probability of the liquid and vapour states. (**B**) Posterior distribution of electric field coupling parameter, *m*, for each combination of water model and conformational state. (**C**) Posterior distribution of intrinsic electric field, $E_{int}$. Since this parameter is independent of the water model, only one kernel density estimate of the distribution is shown for each conformational state of the 5-$HT_3$ receptor.

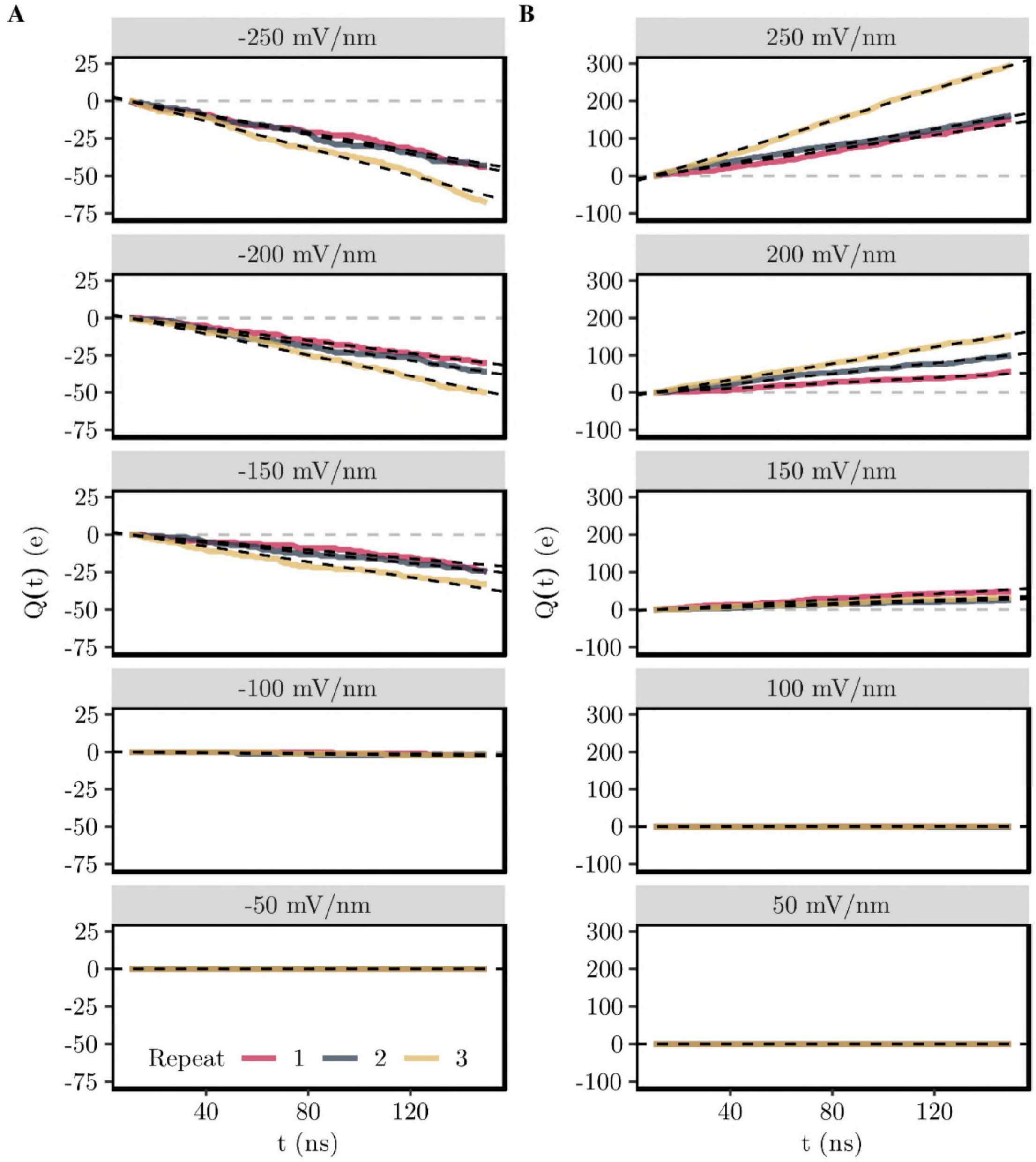


*Figure S4:*

Cumulative charge flow through the M2 helix bundle of the closed state 5-$HT_3$ receptor (PDB ID: 4PIR) due to $Na^+$ ions. Coloured lines indicate the charge flow measured during three independent 150 ns long MD simulations employing the TIP3P water model, while dashed black lines represent a linear fit to these data. The dashed grey lines serve as reference and indicate the case of no electric current. (**A**) Charge flow under the influence of a negative electric field ($E < 0$). Positive charges are driven from the extracellular space to the intracellular domain. (**B**) Charge flow under the influence of a positive electric field ($E > 0$). The direction of charge flow is reversed here.

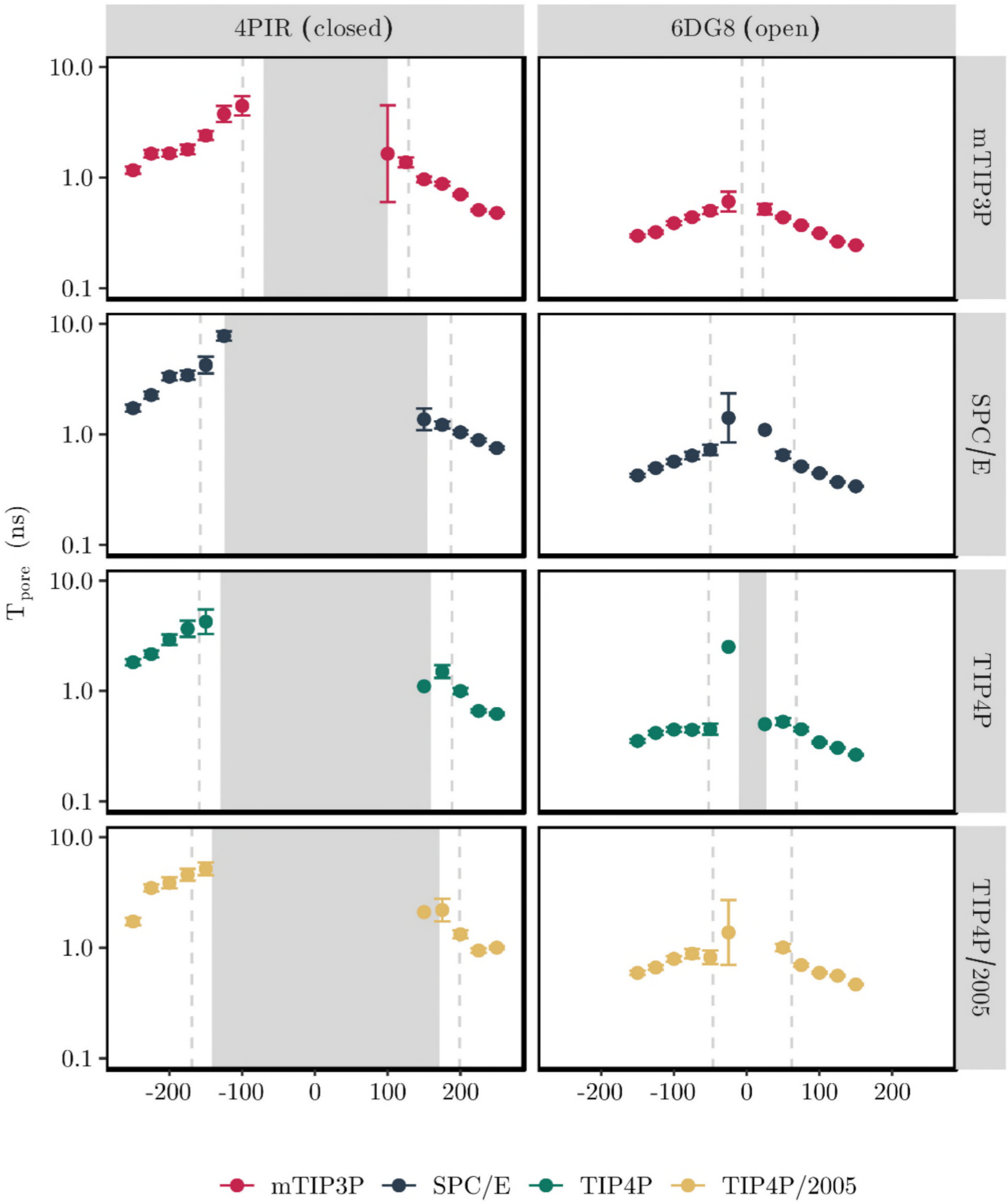


*Figure S5:*

Dwell time of ions inside the M2 helix nanopore as a function of *E*-field strength. Background shading indicates the regime in which the channel is de-wetted, while the dashed lines mark the transition to a mostly hydrated channel. Ions that exited the pore in the direction from which they entered are not included and no distinction is made between $Na^+$ and $Cl^-$ ions.
